# Correcting Chimeric Crosstalk in Single Cell RNA-seq Experiments

**DOI:** 10.1101/093237

**Authors:** Atray Dixit

## Abstract

As part of the process of preparing sequencing libraries that include unique molecular identifiers (UMIs) such as many single cell RNA-seq (scRNA-seq) libraries, a diverse template must be amplified. During amplification, spurious chimeric molecules can be formed between molecules originating in different cells. While several computational and experimental strategies have been suggested to mitigate the impact of chimeric molecules, suitable approaches for scRNA-seq experiments do not exist. We demonstrate that chimeras become increasingly problematic as samples are sequenced deeply and propose both supervised and unsupervised computational solutions. These solutions are validated in the context of a deeply sequenced species mixing experiment, and, orthogonally, using replicate PCR amplifications of the same scRNA-seq library. Our code is publicly available at https://github.com/asncd/schimera.

## Main

By genomically profiling single cells, researchers can evaluate the extent to which cellular variation impacts physiology. As innovations in barcoding and microfluidics have enabled thousands of cells to be processed in a single pot, there have been concomitant reductions in the time and cost required to profile a single cell^1–6^. While pooled library generation has its advantages, it can also create artifacts. Specifically, during PCR, either for library amplification or to selectively enrich a subset of molecules of interest from a complex RNA library^4,7–9^, chimeric molecules can form. These chimeras are formed between transcripts originating in one cell and the barcode corresponding to another (**Figure 1A**).

**Figure 1:**
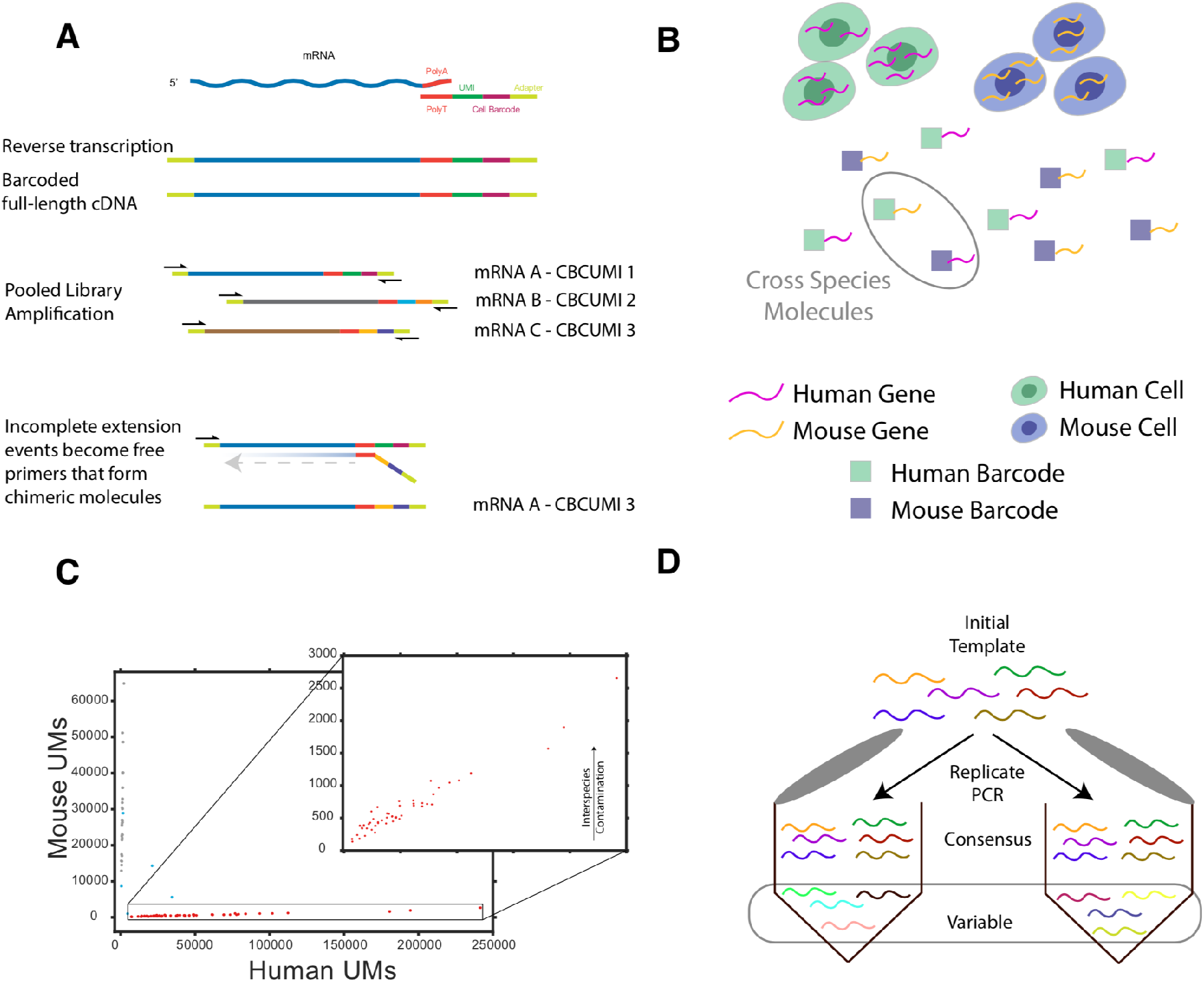
The formation of chimeric molecules during scRNA-seq and evaluation approaches. (**A**) An overview of a scRNA-seq library construction protocol in which RNA capture and reverse transcription is followed by pooled library amplification using PCR Incomplete extension events or template switching can form chimeric molecules. (**B**) Cross species molecules can serve as supervised labels if PCA chimeras are a major generative source for them. (**C**) A scatter plot of cell barcodes and their abundance of Human UMs (x-axis) and Mouse UMs (y-axis). The enlarged subpanel shows the interspecies contamination of mouse UMs is correlated with the abundance of Human UMs. (**D**) Molecules occurring inconsistently across deeply sequenced PCR replicates are likely to be chimeric events and serve as supervised labels for assessing a classifier.

We discuss two cases in which chimera formation can be evaluated: 1) Experiments in which several distinct cell states are present, from the species mixing “barnyard” experiments in which human and mouse cells are mixed together before being processed (**Figures 1B, C**), to the more general case of distinct expression clusters within a species and 2) replicate PCR amplifications (**Figure 1D**). We demonstrate that in both cases contaminating molecules tend to possess distinct characteristics. As such, it is possible to identify and computationally filter them from subsequent analysis.

In a deeply sequenced a species mixing experiment collected using Drop-seq^2^, many cross-species molecules were observed (**Figure 1C, Table S1**). These molecules can come from 1) ambient RNA, 2) barcode collisions due to random chance, and 3) chimeric molecules formed during PCR. We find that 1.7% of all unique molecules (UMs) in cells unambiguously assigned to one of the two species (removing obvious doublet cells) came from the opposite species. If, to obtain an upper bound, we assume all of these molecules to come from chimeric events, then, including presumed human-human and mouse-mouse chimeras, up to 5.2% of all molecules considered in this experiment would be chimeric (based on expected fraction of 33% coming from cross-species products of **Table S2**).

Cross-species UMs are associated with a lower number of reads per molecule. As such, they increase in relative abundance as a function of sequencing depth (**Figure 2A**). Paradoxically, sequencing deeply can reduce the statistical power to detect differential expression.

**Figure 2:**
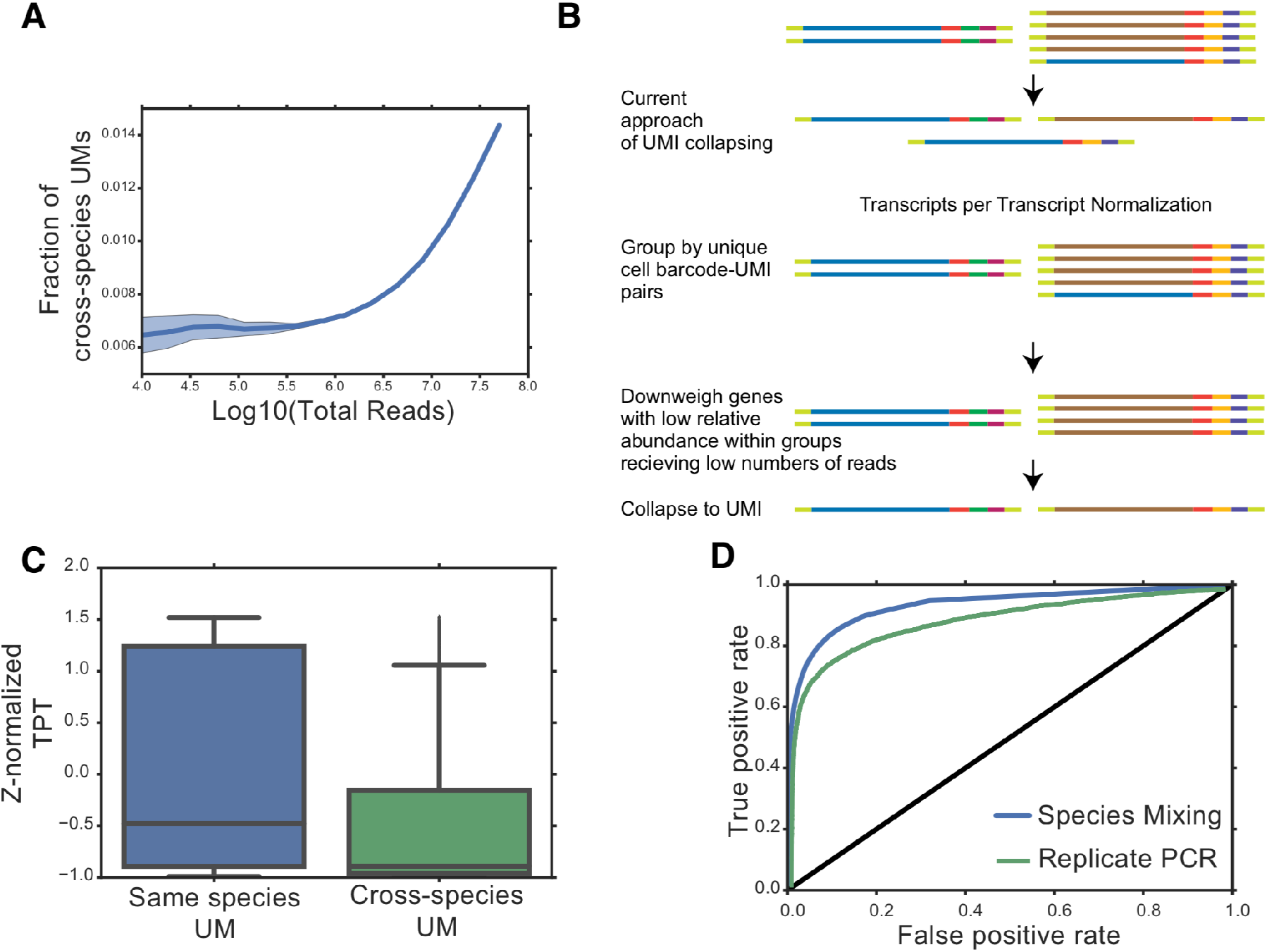
Strategies for Correcting for Chimeric Molecules. (**A**) Rarefaction analysis shows the fraction of cross-species molecules observed increases with sequencing depth. Shaded error region is 1 s.d from 10 independent subsampling analyses. (**B**) Current scRNA-seq quantification using UMIs collapses molecules into unit counts regardless of overall read abundance. An alternative approach is suggested in which molecules are first grouped by cell barcode-UMI pairs and then molecules that have a low relative abundance are filtered before collapsing. (**C**) Boxplots showing how the TPT distribution (Z-normalized within each species) varies with label. (**D**) Receiver operating characteristic curves for a random forest classifier for the deeply sequenced species mixing Drop-seq experiment (blue) and replicate PCR (green).

Depending on the diversity of the UMI barcodes used in library preparation, most pairs of cell barcodes and UMIs should be uniquely associated with a single gene, but we observed far more genes per cell barcode UMI tag in the species mixing experiment than expected based on the observed barcode diversity (**Figure S1C**). PCR chimeras can cause swapping between cell specific barcodes (**Figure 1A**), and are more likely to happen during later stages of PCR. Resultantly, the chimeric molecules will have a lower number of reads per molecule and are likely to have another gene containing the same cell barcode-UMI pair containing a higher number of reads. We introduce a notion of transcript per transcript (TPT) as an additional measure of a chimera formation (**Figure 2B**). In the deeply sequenced Drop-seq library, cross-species molecules are strongly associated with lower TPT (**Figure 2C**) as well as fewer reads per UMI (**Figure S1B**). These features of cross-species molecules were not strongly correlated with one another (**Figure S1F**) and we were able to increase the power to detect differential expression between human and mouse cells by filtering molecules with a TPT less than 0.02 (**Figure S1E**). We found 10% of molecules have a TPT less than 0.01 and almost 30% have a TPT less than 0.05 (**Figure S1D**). We observed far more genes per CBC-UMI tag than we would expect based on the diversity of UMI barcodes present (**Figure S1C**). Our unsupervised filtering approach consists of a simple thresholding step to remove molecules with a TPT and/or number of reads less than a value that can be determined by observing the distribution of these two variables on a dataset specific basis (for example a threshold between 1 and 4 reads per molecule may be warranted based in the Drop-seq data after observing the distribution (as seen in **Figure S1A**).

We develop an alternative supervised approach that relies on the presence of distinct cell states in the experiment. A subset of genes that should be mutually exclusive between the cell types (for example the most extremely differentially expressed genes) is then used to create class labels. For example, in the species mixing experiment, cross-species molecules can be clearly identified. We train a random forest classifier on the species mixing experiment using the following features of each UM (**Figure 2D**). The classifier’s AUC is 0.93. However, we note that our performance evaluation is slightly contaminated by the rate of human-human and mouse-mouse chimeras that should be present (contributing an unfair number events labeled as “false positives”). Specifically, the false positive rate is defined as:

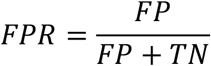

There are 3,394,611 molecules that are not cross species chimeras (our labeled set of “negatives”). Of those, we estimate on the order of 120,000 molecules are actually within-species chimeras. As such, if all chimeras are detected, including the within-species chimeras, the FPR will be at least 0.034. A rough approximation of the max AUC is 0.98 (subtracting the triangle defined by an FPR of 0.034).

By setting the acceptable FPR to 0.034, 4.55% of all molecules are filtered (from before, an estimated 5.2% are chimeras).

In a more shallowly sequenced 10X genomics sample available at - http://support.10xgenomics.com/single-cell/datasets/hgmm_1k that was sequenced to a depth of 69,793 reads per cell and an approximate saturation of 24.4%, we still noted significant cross-species contamination that could be classified somewhat accurately (**Figure S2A,B)**.

Finally, we orthogonally evaluated our unsupervised metrics and supervised approach on data obtained by enriching for a subset of cell barcodes from a transcriptome library as described previously^10^. The enrichment was performed in triplicate. We found that molecules that consistently appeared across PCR replicates differed from those occurring in only one replicate based on TPT and reads per molecule. A supervised classifier combining was able to obtain an AUC of 0.90.

PCR chimeras in scRNA-seq can create challenges in interpreting results. The tendency of chimeras to have fewer reads per molecule can create difficulty in obtaining sensitive detection of medium to lowly expressed transcripts while retaining power to detect differential expression between more highly expressed transcripts (as sequencing depth increases to detect lowly expressed transcripts, chimeras amongst highly expressed transcripts are detected). Chimeras can artificially increase estimates of sequencing saturation if low read abundance chimeras are counted. When performing enrichment PCR to focus on a subset of the library, there will be an increased prevalence of the chimera problem due to higher sequence similarity between amplified molecules.

Longer cell barcodes and UMI’s should help with disambiguating random barcode collisions from true chimeras. Droplet PCR, during which input molecules are amplified independently (**Figure S2C**), may also prevent chimera formation and create more uniform amplification. Pipelines that process scRNA-seq should begin to incorporate some level of chimeric correction. While certainly not perfect, we hope that our supervised and unsupervised methods will help tackle these mythically monstrous mixed molecules in scRNA-seq.

## Acknowledgements

The authors thank all members of the Regev Lab for helpful discussions, encouragement, and support, especially Rebecca Herbst for her help with early brainstorming. Carl de Boer and Christoph Muus provided insightful feedback on the manuscript. Navpreet Ranu shared an early version of his dataset. AD was supported by a NDSEG fellowship.

## Statement on competing interest

A.D. is a founder of and equity holder in Coral Genomics.

---

**Figure S1:**
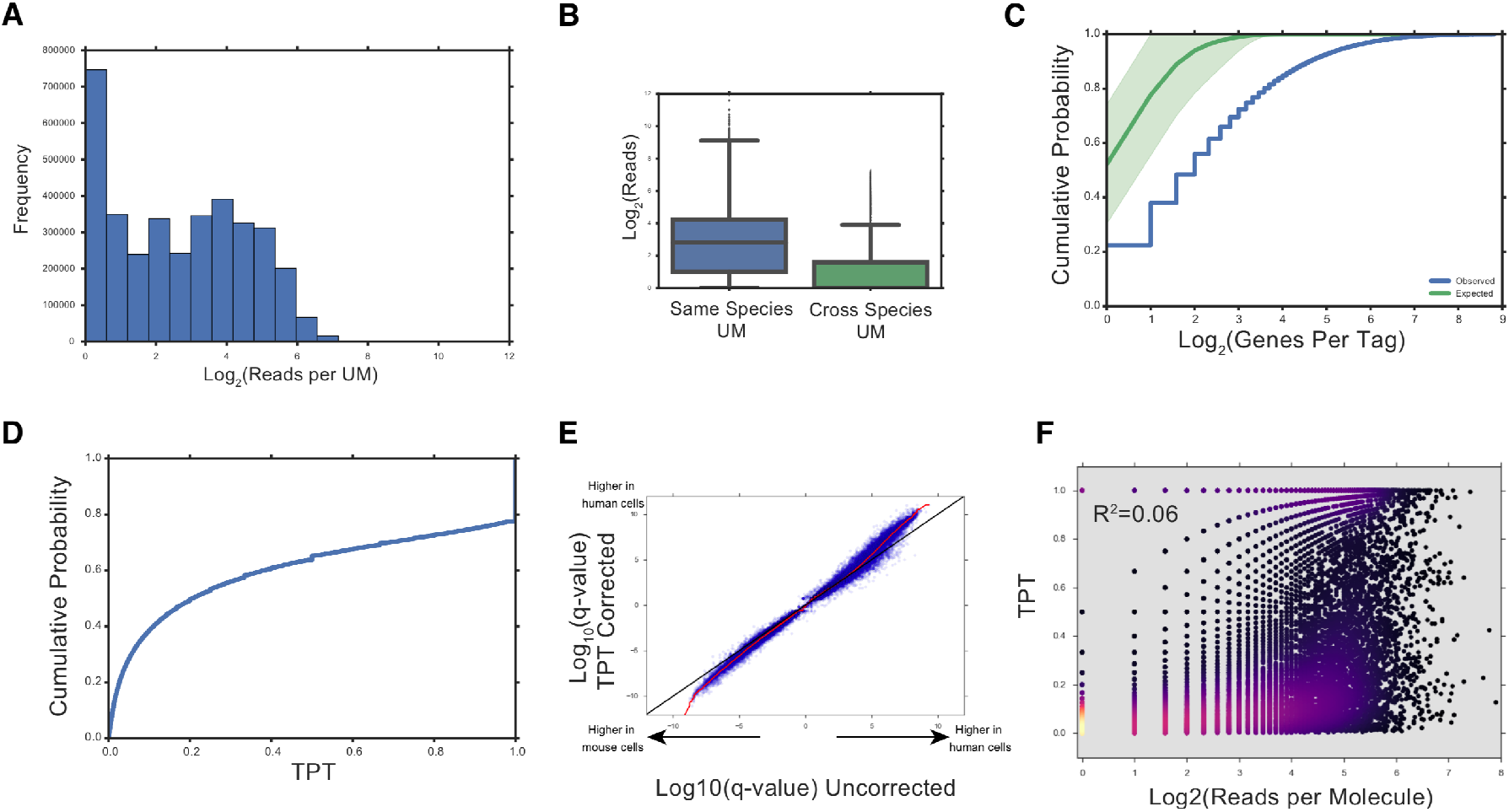
Additional Metrics related to Chimerism. (**A**) Histogram of log_2_(reads per molecule) in this experiment (**B**) Box plots compare reads per molecule for cross species molecules and same species molecules. The box plots denote three quartiles, distribution with whiskers, and outliers as dots. (**C**) Cumulative distribution of the number of genes per unique CBC-UMI tag. Blue line indicates observed distribution. Green line indicates expected distribution based on UMI length and base-pair composition on a per cell basis with 1 s.d shaded in green. (**D**) Cumulative distribution plot of Transcripts per Transcript (TPT) in a deeply sequenced Drop-seq experiment. (**E**) Scatter plot comparing signed log_10_(q-values) for differential expression between human and mouse cells before and after removing molecules with TPT<0.02. Red line is smoothed lowess fit. (**F**) Scatter plot of the relationship between reads per molecule and TPT; the points are colored by local density with black being low density and bright yellow corresponding to high density.

**Figure S2:**
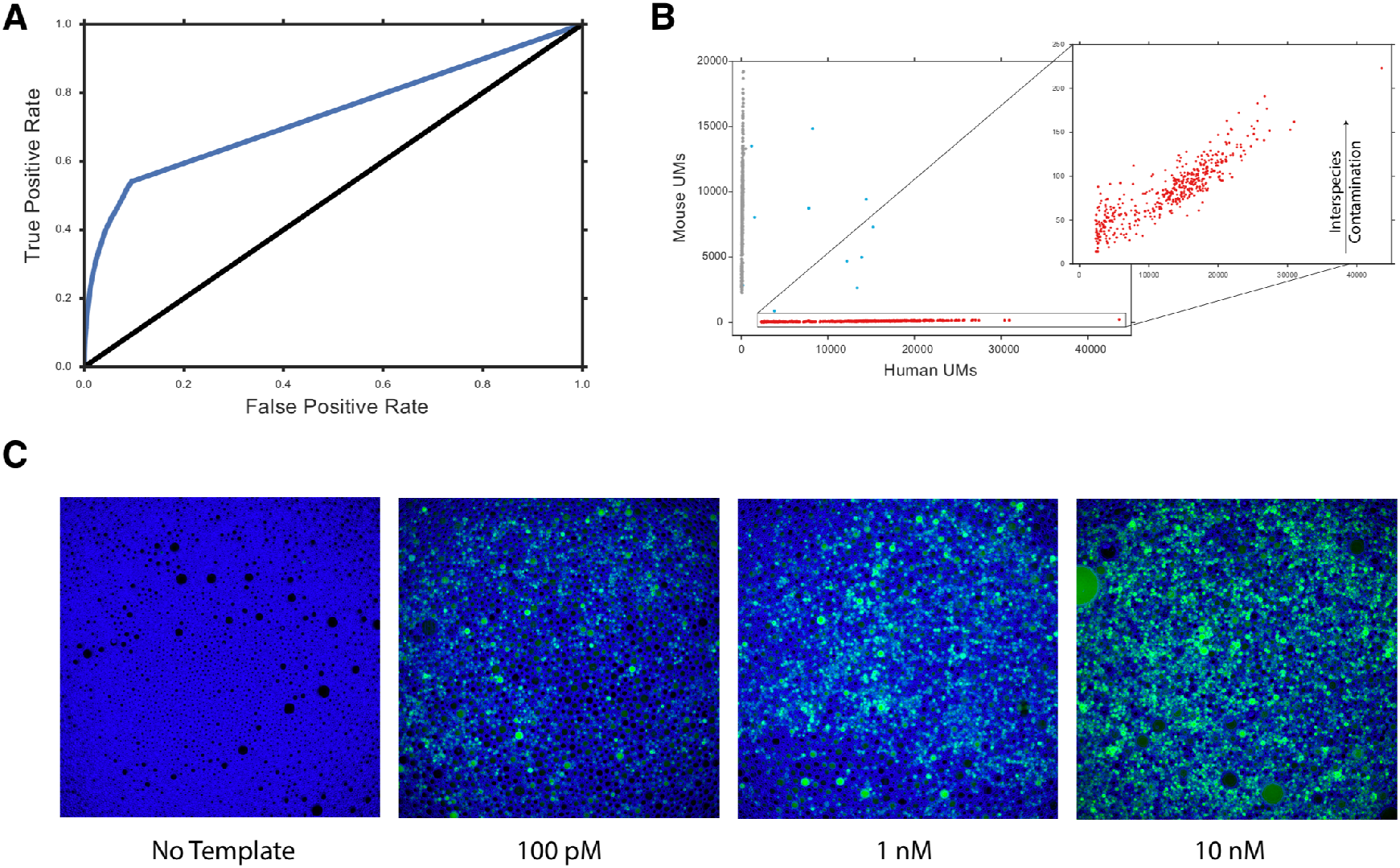
Future Applications, 10X and droplet PCR. (**A**) Receiver operating characteristic curves for a random forest classifier applied a shallowly sequenced (24% duplication rate) 10X dataset. (**B**) A scatter plot of cell barcodes and their abundance of Human UMs (x-axis) and Mouse UMs (y-axis). The enlarged subpanel shows the interspecies contamination of mouse UMs is correlated with the abundance of Human UMs. (**C**) Fluorescent imaging of vortex emulsion PCR droplets containing increasing amounts of a diverse template. SYBR green was added to the PCR mix. Blue outlines were obtained from the bright field image of the droplets.

## Methods

### TPT Normalization

For transcript *i* in a set of molecules that share a unique pair of cell barcode and UMI, *u*, TPT is defined as:

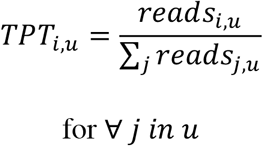

This metric is equal to 1 if a particular cell barcode and UMI pair is unique in the dataset and will be small for a molecule that has a low proportional share of all reads for a particular cell barcode UMI pair.

This normalization approach is applied to the “*molBC*.*txt*.*gz*” in the case of Drop-seq libraries and the *molecules_info*.*h5* file in the case of 10x libraries to filter out molecules with a low TPT. The full gene expression matrix collapsed to UMIs is reconstructed from the filtered molecule set for downstream analysis.

### Downsampling analysis

In order to perform downsampling analysis, the *molBC*.*txt*.*gz* from a Drop-seq species experiment was used. Sampling without replacement was performed on the molecules weighted by their read counts to create downsampled versions of the *molBC*.*txt*.*gz* file. The percentage of cross-species UM was computed as a function of the level of downsampling.

### Supervised Classifier Approach

The following features were considered for the supervised classifier:

- Log_2_(reads per UM); Unique molecules with low number of reads are more likely to be chimeric events.
- TPT; Low TPT is more likely to be a chimera
- Log_2_(total mRNA abundance)
- Log_2_(total CBC abundance)
- Log_2_(Gene Length); Longer genes are expected to be more likely to form chimeras
- GC content; High GC content is also expected to be more likely to form chimeras

We found that reads per UM on its own is able to provide more than half of the overall classification AUC in most cases. A random forest classifier from sklearn in python was used to perform the classification: *sklearn*.*ensemble*.*RandomForestClassifier(n_estimators=300,oob_score=True,n_jobs=-1,class_weight=‘auto’,verbose=1)*

### Replicate PCR

10X libraries were generated following the manufacturer’s instructions. The sheared and ligated products were then amplified using two primers: one targeted to specific cell barcodes and another targeting the Ilumina P7 flowcell adapter on the opposite end.

Full code available at https://github.com/asncd/schimera

**Table S1:**
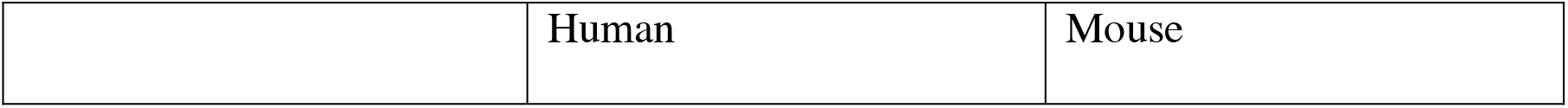
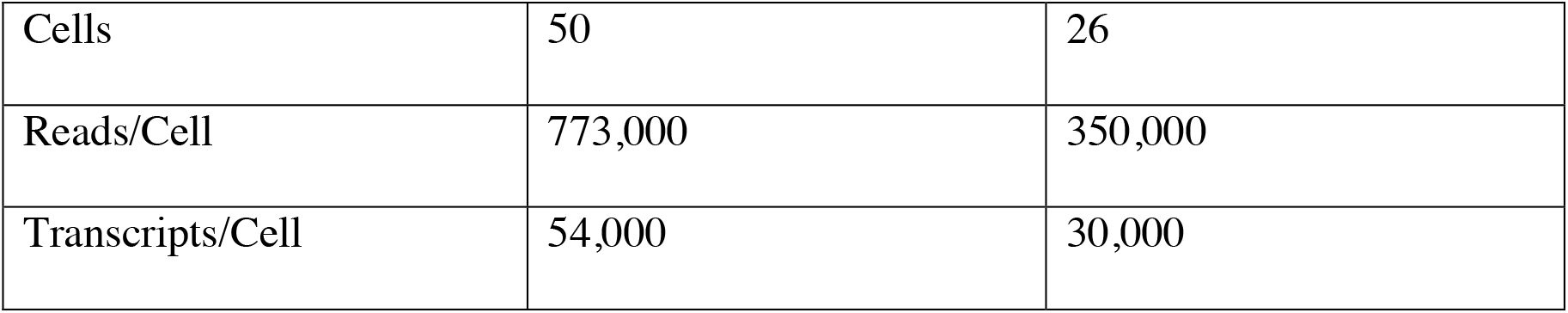
Experimental Summary. Overall cDNA duplication rate (percent of molecules with more than 1 read = 79%)

**Table S2:** Expected relative chimera rates assuming independence (observed numbers of events shown in parentheses, expected number of events shown in brackets).

|  |  |  |
| --- | --- | --- |
| *Expected Relative Chimera Rate* | **Human Cell** | **Mouse Cell** |
| **Human Gene** | 0.63 [n=113,000] | 0.15 (n=27,230) |
| **Mouse Gene** | 0.18 (n=32,014) | 0.04 [n=7,200] |

## Notes

### Competing Interest Statement

The authors have declared no competing interest.

### Summary of Updates

inclusion of species mixing experiment

